# Habitat affinity and density-dependent movement as indicators of fish habitat restoration efficacy

**DOI:** 10.1101/712935

**Authors:** Carlos M. Polivka

## Abstract

Conceptual and methodological tools from behavioral ecology can inform studies of habitat quality and their potential for evaluating habitat restoration in conservation efforts is explored here. Such approaches provide mechanistic detail in understanding the relationship between organisms and their habitats and are thus more informative than correlations between density and habitat characteristics. Several Pacific salmon species have been the target of habitat restoration efforts for the past 2-3 decades, but most post-restoration effectiveness studies have been limited to correlative data described above. In mark-recapture assays from four different study years, the affinity of sub-yearling Chinook salmon (*Oncorhynchus tschawytscha*) and steelhead (*O. mykiss*) for stream pools restored with or created by engineered log structures was greater than that for pools without restoration, though with high interannual variability. From corresponding distribution and density data, it was clear that habitat affinity data are not always concordant with single observations of density. The same was true of the correlation between either affinity or density and physical characteristics of pools, although depth and current velocity had some explanatory power for both responses in Chinook. Movement into pools by Chinook during the assays indicated that restored pools can support more immigrants at a given density than can unrestored pools; however no such pattern emerged for steelhead. Variation among individuals in body condition has implications for population-wide fitness, and such low variation was correlated with stronger affinity for pools in Chinook regardless of restoration status. This suggests that pools may mediate habitat-related trade-offs and that restoring them might have a positive effect on fitness. Thus affinity, immigration, and condition data give much-needed mechanistic indication of habitat selection for restored habitat via an apparent capacity increase and those potential fitness benefits. This is stronger support for restoration effectiveness than density differences alone because density data 1) may simply indicate redistribution of fish from poor to good habitats and 2) are not adequate to show correlations between restoration and positive change in traits correlated with fitness.

## Introduction

Structures placed in streams to create or augment pools are a significant part of restoration efforts in the Pacific Northwest (Roni et al. 2002) because these habitats are important to the rearing phase of the life cycle of salmonids (Roni et al. 2008). For conservation agencies to evalute effectiveness of restoration efforts, appropriate metrics are required (Block et al. 2001). However, such studies usually rely upon comparisons of distribution and density of individuals among both restored and unrestored habitat (Bond and Lake 2003, Roni et al. 2008, Hillman et al. 2016). This is an important basis for evaluation, but much of the literature shows small or no effects of restoration when only distribution and density are considered (Roni et al. 2008, Whiteway et al. 2010, Stranko et al. 2012). Furthermore, the inferential power of the results is limited by inadequate replication of structures and/or observational scale mismatched with treatment scale (Roni et al. 2002, McMillan et al. 2013, Freedman et al. 2016, Polivka et al. 2019), limiting the ability to detect and quantify the seasonal, annual, and among-species distribution patterns (Bradford and Higgins 2001). This does not necessarily indicate poorly placed or targeted restoration activities, but rather the need for consideration of more robust metrics, including those that more directly describe habitat selection (Conrad et al. 2011, Kotler et al. 2016). These metrics include: 1) site fidelity, 2) density dependent movement, and 3) dependence of traits correlated with fitness, like body condition, on habitat selection related movement, all of which can be studied with relatively simple mark-recapture assays.

Site fidelity describes affinity to a habitat unit, in which individuals maintain territories and/or to which they return after life-history-related movements or some other displacement (Greenwood 1980, Merkle et al. 2014). This behavior is presumably driven by habitat effects on fitness (e.g., growth, survival) that can vary at different spatial and temporal scales (Switzer 1993). Affinity of fish to relatively small activity centers, sometimes even as small as a single stream pool, can be identified by mark-recapture studies (Borkholder et al. 2002). Mark-recapture studies in stream salmonids have shown how movement and site fidelity of individuals can vary widely at the site or reach scales (Kahler et al. 2001, Sogard et al. 2009, Myrvold and Kennedy 2016). Movement at the scale of habitat units (e.g., stream pools), however, can be independent of reach-scale movement (Rodríguez 2002), again emphasizing the need for appropriately scaled studies when identifying habitat selection patterns among restored and unrestored habitats.

Density dependence may determine the capacity of a habitat for further settlement and depends on the current occupancy of that habitat. In behavioral ecology, ideal free distribution theory (Fretwell and Lucas 1970, Kennedy and Gray 1993, Houston 2008) describes the density dependent settlement of unoccupied habitat. At low density, there is movement of individuals as they sample the mosaic of habitat patches for the optimal one and attraction to that habitat may be low. As settlement of habitats proceeds, under IFD assumptions, better habitat can support increasing levels of immigration and settlement until it reaches capacity at some optimal density. Then, at higher density, individuals are less likely to immigrate to or remain in the habitat due to crowding; thus immigration will decrease again and emigration will increase (Morris 1988). Observations of density-dependent immigration into a habitat unit (or emigration out of habitat units) can therefore indicate differences in habitat capacity (Gundersen et al. 2002, Rémy et al. 2014). For a given density of individuals occupying the patch, better habitat will support more immigrants.

Resource-driven habitat affinity at the habitat unit scale for fish can involve features such as food availability and cover from predation risk (Giannico and Healey 1999), both of which can be affected by in-stream habitat restoration. Foraging opportunities and cover are complementary resources that often contrast among habitat patches, such that individuals move according to the current levels of risk and food availability (McNamara and Houston 1987, Cresswell 1998, Brown and Kotler 2004). Under these circumstances, the amount of risk a forager will assume can both depend on and affect that individual’s physiological condition (McNamara and Houston 1990, Houston et al. 1993, Kotler et al. 2010). Empirical evidence of such state-dependent habitat selection behavior can provide important clues about habitat quality (Olsson et al. 2002) and thus potentially inform restoration efforts.

The “asset protection principle” (Clark 1994) predicts that individuals in reduced condition take greater risks to obtain food, whereas individuals in good condition remain in safe patches, taking few foraging risks. Under asset protection, those individuals that start in low condition and take risks subsequently increase in condition at a faster rate than individuals that start in high condition and take fewer foraging risks (Luttbeg and Sih 2010). In the short term, this maintains multiple behavioral strategies manifested within a population (Wolf et al. 2007). However, individuals start to converge in their relative condition and measured variation among them will be reduced because most individuals are in moderate-to-high condition and will require similar levels of caution and vigilance (van Gils et al. 2008). Fitness may be optimized or maximized at this point given that further attempts to increase condition would involve additional foraging and thus additional exposure to predation risk (Clark 1994, Clark and Ekman 1995, Nonacs 2001). There are alternative behaviors, however, that result in further divergence among individuals in condition if good condition emboldens those individuals to take more risks, whereas poor condition results in more time and energy allocated to vigilance, predator detection, and thus further decrease in condition with missed foraging (“state-dependent safety;” Luttbeg and Sih 2010, Sih et al. 2015). The foraging and predation avoidance strategy employed (asset protection vs. state-dependent safety) can also be modulated by resource availability and relative predation risk, with highly variable responses observed (Sih et al. 2015).

There are relatively few empirical examples that show the fitness implications of asset protection behavior (Sinclair and Arcese 1995, Lind and Cresswell 2005), despite the fact that growth and condition-related traits are usually correlated with fitness including in salmonid fishes (Zabel and Achord 2004, Ebersole et al. 2006, Tucker et al. 2016). Empirical evidence of asset protection generally shows that foraging effort decreases and predator avoidance activity (e.g., vigilance, apprehension, exposure avoidance) increases with increasing energetic state of foragers (Olsson et al. 2002, van Gils et al. 2008, Matassa and Trussell 2014). Experimental manipulations of forager state, food availability and predator presence/exposure can even quantify the different levels of risk-taking under asset protection (Kotler et al. 2004, 2010). Alofs and Polivka (2004) and Polivka (2007) used this method to show that some estuarine fishes reduce foraging in response to predation and that associated among-individual variation in growth and/or condition indicated that this was likely asset-protection behavior (Polivka 2011).

In streams, shallower, faster current velocity habitats offer rapid delivery of drifting aquatic macroinvertebrates as food, but higher risk of predation, particularly by avian predators. Deeper pools created by log structures, on the other hand, often offer cover from predators, but slower delivery of food. The energetic cost of swimming in the faster flowing, high food delivery habitat is a factor in how much energy may be obtained (Piccolo et al. 2008) and further underscores how pools offer the opportunity for asset protection. Individual based models have made predictions of movement between these complementary habitat types (Railsback et al. 1999, Railsback and Harvey 2002), and there is both laboratory (Reinhardt and Healey 1999) and conceptual (Reinhardt 2002, Conrad et al. 2011) support for asset protection behavior in stream salmonids. Empirical evidence is more difficult to obtain in a field setting (Bradford and Higgins 2001), and sometimes the association between the predation risk regime and the predicted reduction in among-individual variation in condition must be first inferred from correlative data (Polivka 2011) and then confirmed with more specific field observations (Polivka et al. 2013) or experiments. Increasingly consistent use of pools in habitat selection may offer the opportunity for asset protection, detectable as reduced condition variability. Higher pool affinity (independent of restoration status), correlated with lower condition variability among pool residents, would be possible evidence of the role of pools in asset protection behavior of fish. However, this may be difficult to distinguish from pool residency that optimizes energetic intake relative to riffles (Rosenfeld and Boss 2001), for example, which nevertheless may be a form of asset protection, given that fitness correlates such as growth rate can converge around a mean associated with relative energy intake (Girard et al. 2004). Whether pools enhance safety or optimize energy, augmenting them with restoration structures, and/or increasing pool frequency by adding structures, may result in asset protection related fitness benefits to fish compared with stream habitat with fewer, smaller, or lower quality pools.

In this multi-year study, I used mark-recapture assays over a short (24-hr) period to ask whether sub-yearling Chinook salmon (*Oncorhynchus tschawytscha*) and steelhead (*O. mykiss*) have stronger affinity for restored habitat compared with unrestored habitat. I tested whether these affinity patterns were identifiable in individual years, and for all years combined to compare among-year variability with the multi-year trend because longer-term data can add rigor to the evidence for observed post-restoration responses. The second question was whether mark-recapture data supported observations of relative density between habitat types, or whether observational data were less robust than is assumed by post-restoration monitoring programs. Lack of concordance, i.e., years in which there is greater density in restored habitat, but no discernible affinity difference, would suggest that simply collecting distribution and density data is insufficient to address mechanisms behind habitat selection patterns. From associated movement data, I examined density-dependent immigration to determine whether restored pools supported more immigrants than unrestored pools across the range of density of fish occupying the pool from one capture event to the next. I calculated a condition index from size data on each individual and found a correlation between variation in body condition among individuals and affinity for pools in Chinook, as well as other trends. Although I could not distinguish between predator-mediated asset protection and other energetic tradeoffs, restoration may have positive effects on fitness through reduction in among-individual variation. The key finding of this approach is that extending restoration effectiveness studies beyond relative density observations can uncover some of the mechanistic detail needed to better understand fish response to the restoration.

## Methods

### Study System

The Entiat River is a major tributary sub-basin of the Interior Columbia River Basin in north central Washington State, USA. Its confluence with the Columbia River is at 49.657° N, 120.224° W. There, a common habitat restoration action is construction of in-stream structures to create rearing pools for sub-yearling Chinook salmon (listed as endangered) and steelhead (listed as threatened). Restoration is linked with multi-agency monitoring to evaluate its effectiveness (Bennett et al. 2016). Chinook juveniles rear in stream pools and emigrate out of the sub-basin to the mainstem Columbia River if suitable overwintering habitat is not available in tributaries like the Entiat (Hillman et al. 1987), but with most smolt outmigration after one year of freshwater residency. Steelhead can rear in the streams for 1-3 years before outmigration. Predation risk primarily comes from birds (belted kingfisher, *Ceryle alcyon*; great blue heron, *Ardea herodias*) and semi-aquatic mammals (e.g., river otter, *Lontra canadensis*. Larger, predatory fish such as resident and fluvial bull trout (*Salvelinus confluentus*) have been observed in deeper pools created by larger in-stream structures such as channel-spanning weirs, but not in the smaller pools created by the engineered log jams (ELJs) that comprised restoration projects in the river. There is no known study in this system directly quantifying the mortality to sub-yearling Chinook and steelhead from their predators.

For this study, I used two closely situated reaches in the lower geomorphic valley segment (Godaire et al. 2009) of the river ~ 5 km upstream of the Columbia (Figure 1). The restored reach (river km 4.5-4.9) has N = 11 ELJ and rock structures installed in 2008, creating pools with mean area = 32.2, range = 7.6-89.7 m^2^. The unrestored reach (river km 5.2-5.5) has N = 11 natural pools usually formed between small boulders, lacking wood cover, and smaller than the ones created by the structures in the restored reach (mean area = 15.4, range = 3.1-42.6 m^2^). Polivka et al. (2015), Polivka et al. (2019), and Polivka and Claeson (in press) showed that unrestored habitat units in much of the lower geomorphic valley segment of the river basin are statistically indistinguishable in terms of fish density, independent of reach. Thus, the reach association of the restored and unrestored pools is a convention for identification rather than an experimental effect.

**Fig. 1:**
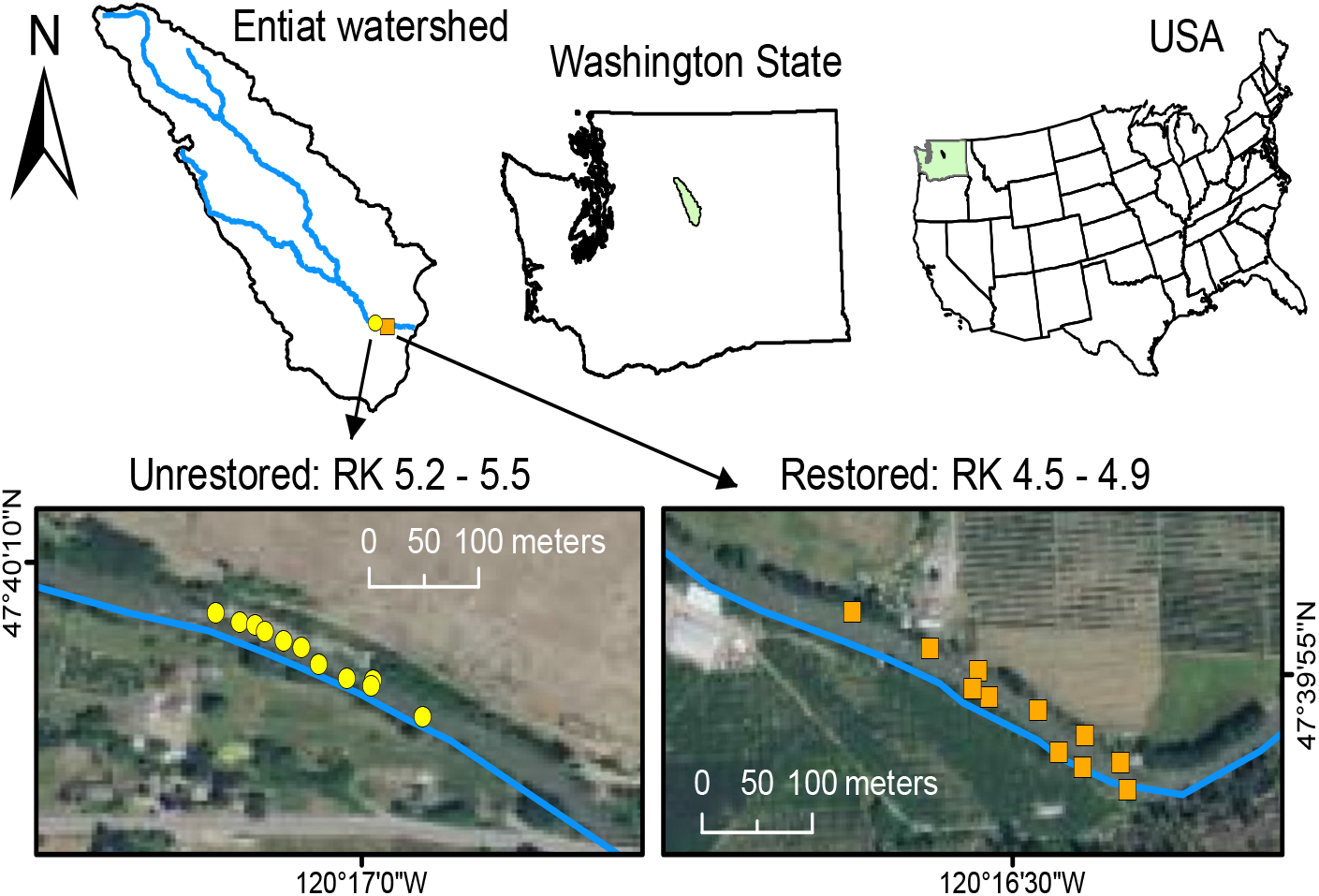
Map showing the location of the Entiat River and the study area with reaches containing restored and unrestored pools (RK = river kilometer, measured upstream from confluence with Columbia River).

### Fish Capture and Marking

All fish handling was conducted under US Dept. of Commerce, NOAA-Fisheries, Permit No. 1422 and is consistent with guidelines published by the American Fisheries Society (Nickum et al. 2004). Field crews conducted behavioral assays during the early rearing season (July) of 2009, 2012, 2013, and 2016; gaps in study years occurred when high river flows affected the schedule of this and concurrent studies (cited above), and, in 2014, owing to a large sediment deposition following a fire in the upper basin.

Electrofishing can be invasive for behavioral studies (Mesa and Schreck 1989); therefore, field crews captured fish using a 3 m × 1.5 m seine with 3 mm mesh. Because pulling a seine along the cobble and rock substrate of this river is ineffective, two field-crew members stood at the downstream end of the pool and held the seine open as two other members snorkeling in the water used large hand nets to coerce fish into the seine and sometimes to capture fish individually using large hand nets. Underwater visibility in the Entiat River is 4-5 m such that two snorkelers can see the entire sampled area and account for any fish, by species, that escaped capture, ensuring high capture success. In unrestored pools, with their smaller size, this meant assurance that capture probability in these pools was 100%, consistent with results reported in Polivka (2010) and Polivka et al. (2015). Across all restored pools, there were < 5 escaped fish total in each year. Fish may also be missed owing to concealment in the submerged part of the ELJ structure, invisible to the snorkelers aiding the seine capture. However, capture of these fish would require electrofishing, which was not tractable because the electric shock stress could affect the relevant behaviors. Therefore, I could not calculate a capture probability by which to adjust the density of pools. Comparison of five years of both seine capture data and snorkel-only fish counts in this study system showed similar density (Polivka et al. 2015), indicating that concealment of fish by ELJ structures does not substantially affect the ability to capture or recapture fish in restored pools. Polivka (2010) and Polivka et al. (in review) obtained data on individual growth by mark and recapture periods ranging from 15-60 d suggesting reliable capture of fish that reside in pools over long periods of time, relative to the time frame of this study. Furthermore, analysis (see below) should give a relatively conservative result if fish indeed have higher affinity for restored pools than unrestored pools.

At each pool, depth, temperature, current velocity, pool area and dissolved oxygen were measured. Captured fish were placed in insulated, aerated buckets and mildly anaesthetized with MS-222 (< 0.1 g · 1^-1^) for 2-3 minutes. Sub-yearling fish in this study system range from 50-75 mm (Chinook) and 35-70 mm (steelhead), depending on growth rates. Following identification and recording of size data (standard length, SL, in mm and mass in g), fish were marked with a subcutaneous injection of visual implant elastomer (VIE; Northwest Marine Inc.). Color combinations and body positions of marks enabled individual identification of fish. Following marking, fish were transferred to another insulated, aerated bucket where they were allowed to fully recover from anaesthetization. The recovery period was at least 10 min, or after a full righting response with fish appearing alert and responsive, before they were released to the capture pool. After 24 hours, the pool was re-sampled and the number of both recaptured individuals and newly captured unmarked fish were noted.

### Data Analysis

#### Affinity patterns over time

Despite reach-scale similarity in fish density, individual pools in this study system vary widely in density whether restored or unrestored (Polivka et al. 2015). This has made comparison of *mean* recapture fractions, and correlation of them with environmental parameters, statistically intractable (Polivka 2010). Across the range of values of fish marked on Day 1 (N_marked_), the number of recaptures on Day 2 (N_recaptured_) should increase linearly. The slope (*β*) of the line fit through the data is an estimate of the average recapture probability for that set of pools. Differences between the slopes (e.g., *β_restored_* > *β_unrestored_*), evaluated by using a N_marked_ × habitat type term in a linear model would indicate differences in affinity for restored and unrestored habitat. Although the ELJ structure can reduce the capture probability to < 1, the primary concern would be marked fish not recaptured due to concealment. However, capture efficiency is assured in unrestored pools; thus, if there are missed recaptures in restored pools a finding of higher affinity for restored pools relative to unrestored pools would be conservative.

I specified generalized linear models (GLMs) for each species, assuming a Poisson error distribution. I assumed *a priori* that the relationship between N_recaptured_ on Day 2 and N_marked_ on Day 1 passed through the origin because pools with no fish captured (or marked) on Day 1 could have no recaptures and were therefore excluded. Parameters tested as predictors were N_marked_, habitat type, and year, with year included or excluded to confirm whether or not annual differences should be tested separately, along with the longer-term trend from combining years. Because pool area is a strong positive correlate of fish density in this system (Polivka et al. 2015), and thus affected the number of fish marked to begin with, it was entered into each model as an offset parameter to prevent fitting a negative value or modeling a trivial positive correlation (Zuur et al. 2009). In this fashion, it helps to rule out observed habitat selection and affinity for restored pools as simply an artifact of restoration creating larger pools. I compared four candidate GLMs for each species to determine the importance of habitat and annual effects: 1) equal slopes of the regression lines for the two habitat types (i.e., no N_marked_ × habitat interaction term) with the year parameter included, 2) equal slopes and no year effect, 3) unique slopes (including interaction term) plus the year effect, and 4) unique slopes and no year effect. I selected the best model using the Akaike Information Criterion (AIC; Burnham and Anderson 2002). If the best model was one of the models that included a N_marked_ × habitat type interaction term, I concluded that fish differed in their affinity for restored vs. unrestored habitat and calculated habitat-specific βs. Given the multi-year nature of the study, I expected the best model to include the year term, justifying within-year analyses. To ensure that the offset parameter (pool area) did not cause some systematic lack of model fit, I re-ran the analyses with pool area designated as a fixed effect.

To address annual differences and to compare affinity with observations of density, I used two sets of GLMs for each year: one for affinity (including the N_marked_ × habitat interaction term) and one to describe fish density on Day 1. I also tested whether physical habitat characteristics (depth, current velocity, temperature, pool area) are consistent predictors of either N_recaptured_ or of Day 1 abundance (N_marked_). A similar analysis of density appears in Polivka et al. (2015), but that study uses repeated data within each year, whereas this analysis uses only data specific to Day 1 of the mark-recapture trials. As such, the density data are comparable to data collected in most restoration effectiveness monitoring efforts in the region (i.e., single observations of fish density; Roni et al. 2015, Hillman et al. 2016).

Selection of the best affinity and abundance GLMs proceeded stepwise with the removal of non-significant (p > 0.05) predictors, until the resultant model contained only significant terms. Model output for GLM in R (R Core Team 2018) provides AIC scores and I used these to confirm that the model with only significant predictors also had the lowest AIC score. These models identified 1) the years in which *β_restored_* and *β_unrestored_* indicated different levels of habitat affinity, i.e., if the best model had a significant interaction term, 2) whether any differences in affinity corresponded to differences among habitats in abundance, and 3) whether affinity and abundance were associated with the same physical attributes of pools.

In all models describing affinity, both for individual years and years combined, I considered potential issues with capture success. On Day 2, captures consisted of N_recaptured_ + unmarked individuals. Unmarked individuals were generally assumed to be immigrants into the pool between Day 1 and Day 2; however, they could also be individuals not captured on Day 1 that remained in the pool. To determine whether this affected model outcome, I re-analyzed all data by assuming that Day 1 individuals observed to have escaped capture remained in the pool as recaptures. Assumptions about any Day 2 individuals that were not captured were too weak to justify further adjustment of the models.

#### Density-dependent immigration, emigration, and capacity

Total immigration likely depends on pool size, so I first examined the linear relationship between pool area and total immigrants with linear regression. To determine whether restored pools allowed for greater density-dependent immigration relative to unrestored pools, I took the number of immigrants (i.e., unmarked fish captured on Day 2) and examined how it was affected by the density of fish maintaining occupancy of the pool (i.e., recaptured) over the 24 hour period. For this relationship, I tested the fit of the data to a Ricker-style function (Ricker 1954). Such functions are of the form *xe^1-x^* and are widely used in fisheries to describe density dependent processes (e.g., Sharma et al. 2005). The specific modification used here is:

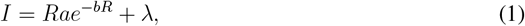

where *I* = number of immigrants, *R* = density of recaptured fish and *a* and *b* describe the shape of the response curve. The peak level of immigration is 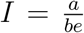 at recapture density 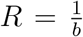 and λ is a term added to represent a constant level of density-independent immigration, particularly given that immigration may be observed at zero density of recaptures. With three parameters in need of estimation (*a*, *b* and λ), data from all years were combined to avoid over-fitting of the model (Anderson 2008). I used total number of immigrants rather than immigrant *density* because pools vary in size and density and two pools of a different size can have the same total density. The larger pool will possibly have the capacity for a greater number of immigrants, but this may translate to low immigrant density for the relative amount of available habitat. Because the total number of pools per year sampled on Day 1 was, at most, 11 and was first reduced by the number of fish in which there were zero captures on Day 1 in any given year, I had insufficient replication to evaluate effects of individual years and individual pools. This is because the error term for each parameter in Equation 1 would require partitioning into habitat, year, and pool effects. This was intractable because the logistical issues listed above resulted in different combinations of pools used each year. Also, different cohorts of fish were sampled each year; thus, each year’s data had a reasonable level of independence and thus were combined as a whole for analysis.

Parameters from Equation (1) for each habitat type (restored or unrestored pools) were estimated by non-linear least squares, which is generally equivalent to maximum likelihood estimation, especially for small sample size when the assumption of normality may not hold (Amemiya 1977, Anh 1988). The output included the 95% confidence interval for each parameter. Differences between habitat types in the parameter value could be identified by non-overlapping confidence intervals, but wide confidence intervals might require an additional test of whether those differences are meaningful. Therefore, I used a randomization procedure to compare the values. I made a random draw from the values in the 95% confidence interval around the parameter estimate and generated a uniformly distributed set of 10,000 values for each parameter in each habitat. From the set of 10,000 values for that parameter, I drew, with replacement, 1000 values, and calculated the difference (e.g., *a_restored_ – a_unrestored_*) for each pair. I then examined the 95% Highest Density Interval (HDI) for the 1000 values of that difference. A parameter was considered different between habitats if the 95% HDIs for the difference did not overlap zero.

Examination of density dependent emigration is a simpler process as emigration is expected to increase linearly with density. However it may also be an artifact of the total number of fish marked in a pool, total pool area, or there may be differences in total emigration by habitat type. Therefore, I constructed another linear model using each of these parameters. To determine whether habitat type affected density dependence, I included a habitat × density interaction term. I performed analysis of variance on these models to identify significant predictors of total emigrants.

#### State-dependent movement, and among-individual variability in condition

To examine whether condition variability among individuals was correlated with habitat affinity as predicted by the asset protection principle, I tested the correlation between the coefficient of variation in the Fulton Condition Index for fish (*K*; Anderson and Neumann 1996) and habitat affinity *β*. Despite its simplicity, I used the Fulton Index because it also offered a rapid, non-invasive way of estimating the body condition of a large number of fish, while minimizing disturbances resulting from handling to the short-term affinity and movements that were under study. The Fulton Index relates length (***L***) and mass (*m*) as:

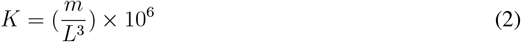

Although the scaling exponent for *L* can vary among species, I used a log(mass) vs. log(length) regression to determine that the exact value was 2.997 ± 0.013 and thus not significantly different from 3. Another critical concern with condition indices such as K that are derived from length and mass relationships is that they can be inaccurate when comparing across age classes and/or when there is sexual dimorphism (Peig and Green 2010). Because this study used only sub-yearlings, a stage at which sexual dimorphism is not apparent, there should not be accuracy problems with *K* in this case. Moreover, the key question is whether among-individual variability in condition is affected by affinity for pool habitat (regardless of restoration status). Here, I used the β values obtained from the linear models which quantified relative differences in affinity for either habitat type in each year. I calculated the coefficient of variation in the condition index (CVCI) among individuals recaptured in each pool.

I used the Pearson correlation *r* to evaluate the correlation between β and CVCI for both species. If pools offer “asset protection” to complement habitats with higher food availability as suggested by Railsback and Harvey (2002) and Bradford and Higgins (2001), then CVCI and the affinity for pools (β) should be negatively correlated. Pool residents (i.e., those recaptured after 24hr) might also show lower variation among individuals than individuals that are immigrating or emigrating. That hypothesis had to be evaluated qualitatively because coefficients of variation summarize large amounts of data and leave too few values from which to generate meaningful statistics. These analyses also do not incorporate direct measurements of predator abundance or any estimate of actual predation risk. Under asset protection, the CVCI would be expected to be greater for fish moving in or out of pools between capture events, i.e., emigrants (those marked on Day 1 and not recaptured on Day 2) or immigrants (unmarked fish captured on Day 2) relative to residents (those recaptured). To consider whether emigration or immigration was simply size-dependent, resulting from larger individuals claiming or holding territories, I compared the mean standard length and CI among individuals from each group with a one-way analysis of variance and post-hoc Tukey HSD tests.

## Results

### Affinity patterns by year

In each year of the study except 2016, 500-700 Chinook and 50-200 steelhead were marked (Appendix Sl: Table 1). In 2016, however, numbers of both species were approximately four-fold lower. In 2016, mean temperature across all habitats was cooler (16.6° C) and mean current velocity was slightly faster (30.3 cm · s^-1^) than in the other three study years. Furthermore those three years did not vary strongly among each other (temperature range = 17.1-18.9° C; current velocity range = 20-27 cm · s^-1^). There was variation in whether restored or unrestored habitat had highest density of either species in the single samples taken for these assays (see below).

**Table 1:** Generalized linear model (GLM) fits of number of recaptures (*N_r_*) to number initially marked (*N_m_*) in 24 hr site fidelity assays in restored vs. unrestored pools (2009, 2012, 2013, 2016). Models considered were: 1) fit lines with equal slopes for both habitats (no *N_m_*× habitat interaction term included) and including study year as a factor, 2) equal slopes and no year effect, 3) unique slopes (with interaction term) and the year effect, 4) unique slopes and no year effect. Model selection by AIC (best fit model in boldface), and habitat affinity indicated where applicable (NA indicates models that do not distinguish habitats by GLM slopes).

| Model | Affinity | ΔAIC |
| --- | --- | --- |
| *a) Chinook salmon* |  |  |
| $N_r = N_m + \text{habitat} + \text{year} + \text{area}$ | NA | 17.5 |
| $N_r = N_m + \text{habitat} + \text{area}$ | NA | 70.7 |
| **$N_r = N_m + \text{habitat} + N_m \times \text{habitat} + \text{year} + \text{area}$** | **restored** | **0** |
| $N_r = N_m + \text{habitat} + N_m \times \text{habitat} + \text{area}$ | restored | 57.2 |
| *b) Steelhead* |  |  |
| $N_r = N_m + \text{habitat} + \text{area}$ | NA | 21.7 |
| $N_r = N_m + \text{habitat} + \text{year} + \text{area}$ | NA | 4.5 |
| **$N_r = N_m + \text{habitat} + N_m \times \text{habitat} + \text{year} + \text{area}$** | **restored** | **0** |
| $N_r = N_m + \text{habitat} + N_m \times \text{habitat} + \text{area}$ | restored | 12.3 |

For both species, the GLMs selected by AIC included the year parameter, and the best model for each contained a significant interaction term (N_marked_ × habitat, P < 0.0001), indicating unique slopes for restored and unrestored pools (Table 1, Figure 2). With all years combined, the affinity of each species was greater for restored habitat; however, for Chinook, the difference was very small (Chinook, *β_restored_* (± se) = 0.258 ± 0.02, *β_unrestored_* = 0.226 ± 0.02; steelhead, *β_restored_* = 0.573 ± 0.04, *β_unrestored_* = 0.388 ± 0.02). There was no indication of a systematic lack of fit with these models, whereas removal of the offset parameter (pool surface area) led to problems with convergence in Model 1 (same slope, no random effect). Thus, the original model specification, with offset parameter, is justified.

**Fig. 2:**
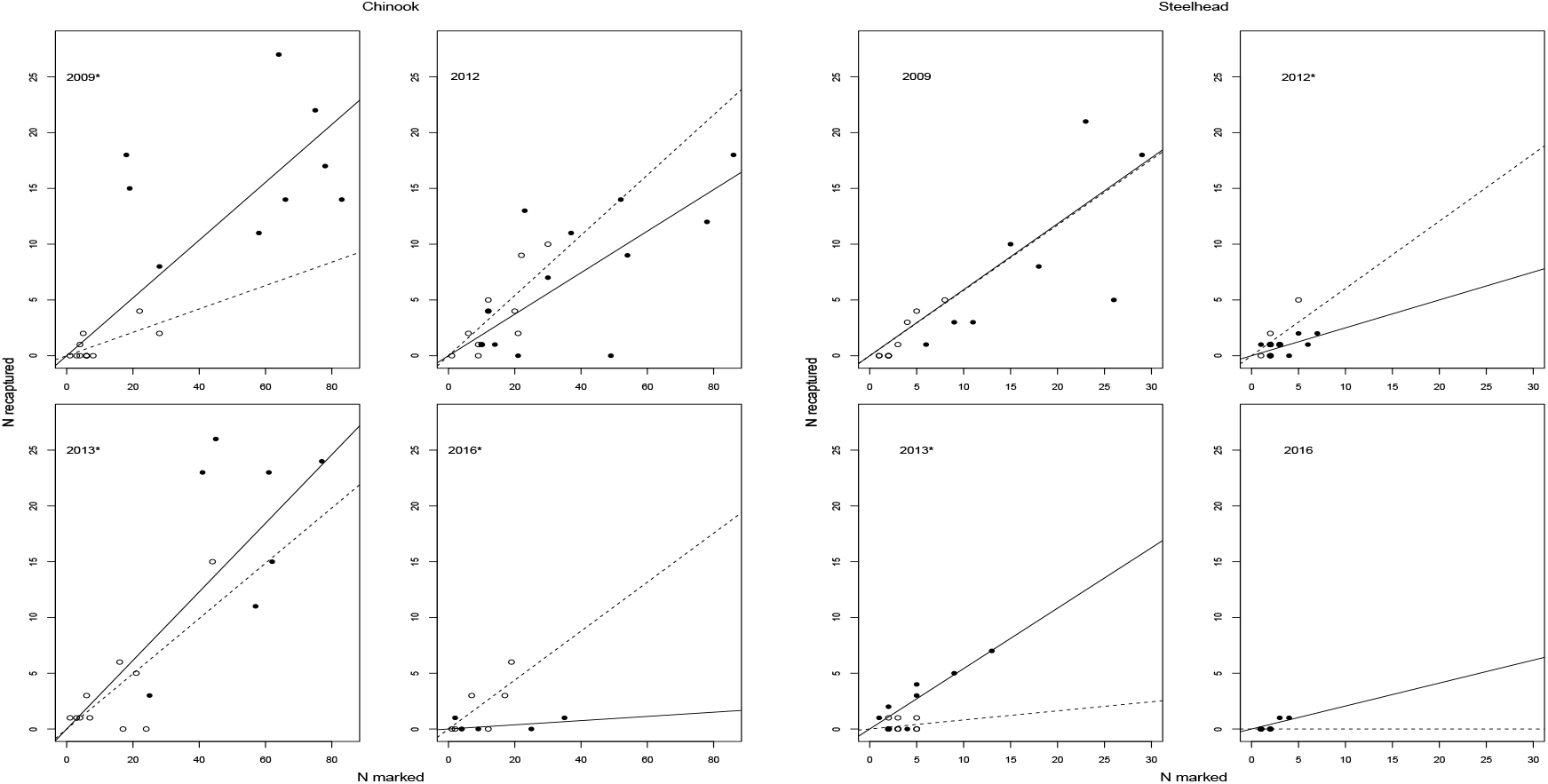
Habitat affinity, shown as the linear fit of *N_recaptured_* vs. *N_marked_*) for restored (solid lines, filled symbols) and unrestored (dashed lines, open symbols) in 24-hr mark-recapture assays in each year for sub-yearling Chinoook salmon and steelhead. Overall habitat differences in affinity for all years combined indicated by GLM fits described in Table 1; slopes of lines and significant within-year differences indicated with * as described in Table 2.

In analyses of individual years, for both species, and for both affinity and density, the model resulting from stepwise selection of terms also had the lowest AIC score, thereby validating stepwise selection. Chinook showed higher affinity for restored pools in 2009 and 2013, but did not differ in habitat affinity in 2012 and marginally favored unrestored habitat in 2016 (N_marked_ × habitat, p = 0.058; Table 2; Figure 2a). Early season habitat affinity and density were concordant only in 2009 and 2013. Affinity was typically correlated with similar pool characteristics as density, including deeper, slower flowing water (Table 2). Steelhead affinity was also variable from year to year (Table 2; Figure 2b) and the strong difference in β with years combined (above) appeared to be primarily influenced by the strong difference favoring restored habitat in 2013 (Table 2). Affinity for unrestored habitat was indicated in 2012; no other year showed a difference in β. Affinity in 2016 could not be determined by GLM because only three unrestored pools had any marked fish (N = 1-2) and, in each case, zero recaptures. Steelhead density favored restored habitats in 2009 and 2013, but not in 2012 or 2016. Physical correlates did not have much explanatory power for either abundance or density in steelhead (Table 2).

**Table 2:**
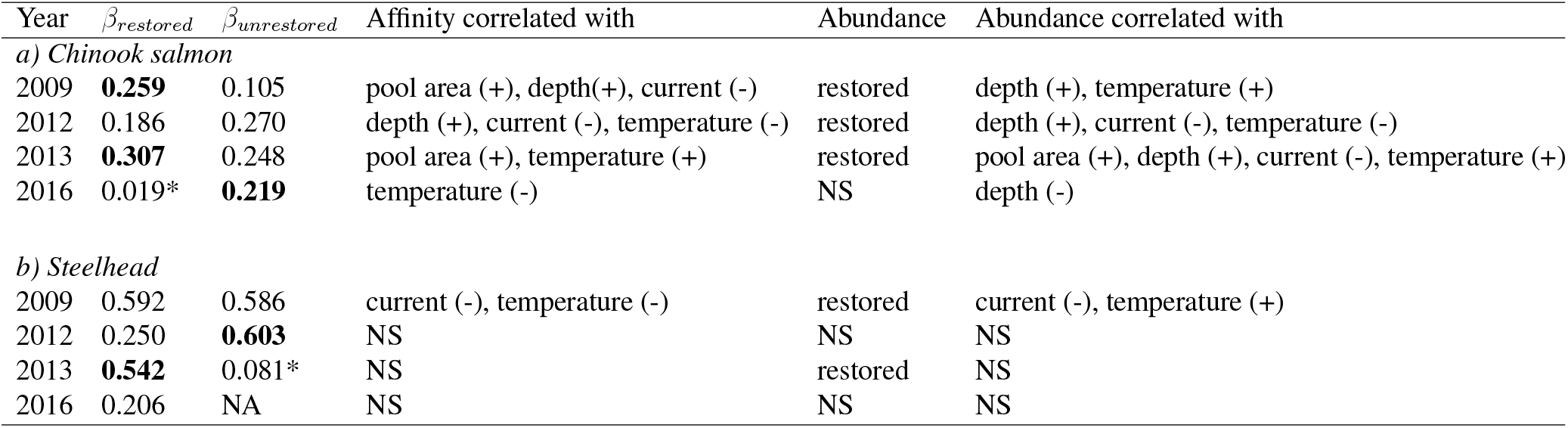
Results of separate GLM analyses of affinity and abundance patterns in a) Chinook salmon and b) steelhead in restored vs. unrestored pools in each year of the mark-recapture assays. The GLM specification for affinity was as in Table 1 (excluding a year term, but including physical habitat covariates) where the N_marked_ × habitat term identified differences in affinity among habitat types. The value of the slope (*β*) is indicated in boldface in habitats with significantly greater affinity. Slopes with * are not different from zero within that habitat type and usually based on low sample size in that habitat × year; NA: Zero recaptures in N=3 pools, only 1-2 fish marked per pool. Abundance differences among restored and unrestored habitats identified by GLMs are indicated. Significant positive (+) and negative (-) correlations of physical habitat parameters shown for each group of models. For both affinity and abundance models, the best model was selected by stepwise removal of non-significant terms.

| Year | $\beta_{\text{restored}}$ | $\beta_{\text{unrestored}}$ | Affinity correlated with | Abundance | Abundance correlated with |
| --- | --- | --- | --- | --- | --- |
| *a) Chinook salmon* |  |  |  |  |  |
| 2009 | **0.259** | 0.105 | pool area (+), depth(+), current (-) | restored | depth (+), temperature (+) |
| 2012 | 0.186 | 0.270 | depth (+), current (-), temperature (-) | restored | depth (+), current (-), temperature (-) |
| 2013 | **0.307** | 0.248 | pool area (+), temperature (+) | restored | pool area (+), depth (+), current (-), temperature (+) |
| 2016 | 0.019* | **0.219** | temperature (-) | NS | depth (-) |
| *b) Steelhead* |  |  |  |  |  |
| 2009 | 0.592 | 0.586 | current (-), temperature (-) | restored | current (-), temperature (+) |
| 2012 | 0.250 | **0.603** | NS | NS | NS |
| 2013 | **0.542** | 0.081* | NS | restored | NS |
| 2016 | 0.206 | NA | NS | NS | NS |

Mean (± se) depth in each habitat type was 56.5 ± 20.1 cm (restored) and 44.8 ± 9.6 cm (unrestored). Mean current velocity was 18.0 ± 10.0 cm · s^-1^ (restored) and 32.8 ± 15.1 cm · s^-1^ (unrestored). Temperature was indicated as a significant correlate in some Chinook models, but the correlation often was opposite in direction for affinity vs. abundance (Table 2). Furthermore, there was essentially no temperature difference between restored and unrestored pools during the assays (Mean temperature, restored pools = 18.01° C; unrestored pools = 18.05° C). Affinity did not seem to track either current velocity or temperature. In 2009, 2012, and 2013, affinity of either species for restored habitat could be strong, moderate or not different from unrestored habitat (Table 2) across the moderate average temperatures or current velocities in those years. In 2016 when average current velocity was faster and average temperature lower (see above), Chinook favored unrestored habitat and steelhead showed no affinity difference.

### Density-dependent immigration/emigration

Regression analysis showed a significant linear increase in the number of immigrants with pool area for Chinook (F_1,71_ = 20.48, p < 0.0001) but not for steelhead (F_1,61_ = 0.634, p = 0.424). The fitted Ricker-style functions for each habitat type (Figure 3) indicated a higher peak Chinook immigration level in restored habitat (*a_restored_* = 115.2, *a_unrestored_* = 27.76) The 95% HDI of this difference among habitats was > zero (range = 3.33-225.74, Figure 4). The confidence intervals for *b* and λ from each habitat overlapped substantially, so they were not analyzed further. The lack of difference in the shape parameter *b* indicates that peak immigration occurs at the same recapture density regardless of habitat type and the lack of a difference in λ indicates that habitats have equal levels of density independent immigration, particularly for pools in which all fish emigrated (zero recapture density). Inspection of a plot of immigrants vs. recaptures revealed no pattern in the steelhead data (Appendix S1: Fig. 1), and an attempt to fit the immigration model resulted in parameters that were not statistically meaningful.

**Fig. 3:**
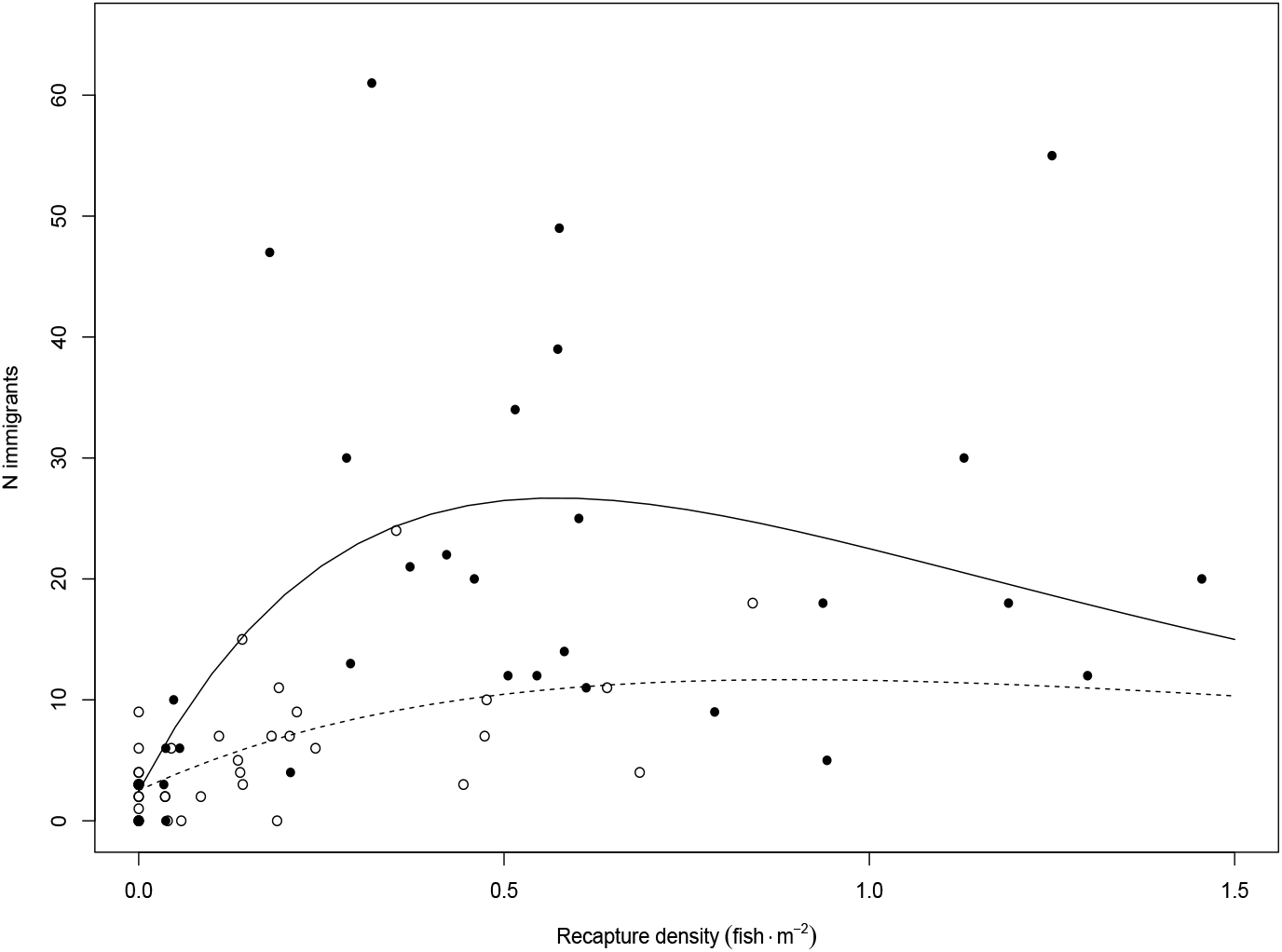
Immigration of young of the year Chinook salmon into restored (solid line, filled circles) and unrestored (dashed line, open circles) pools over a 24-h period as a function of the density of fish maintaining affinity for those pools over that period (recaptures). Density dependence is described by the fit of the Ricker model curve (see Eq. 1) and non-linear least squares estimation of parameters indicates higher total immigration (significantly different *a* term in Eq. 1) in restored pools (see Fig. 4).

**Fig. 4:**
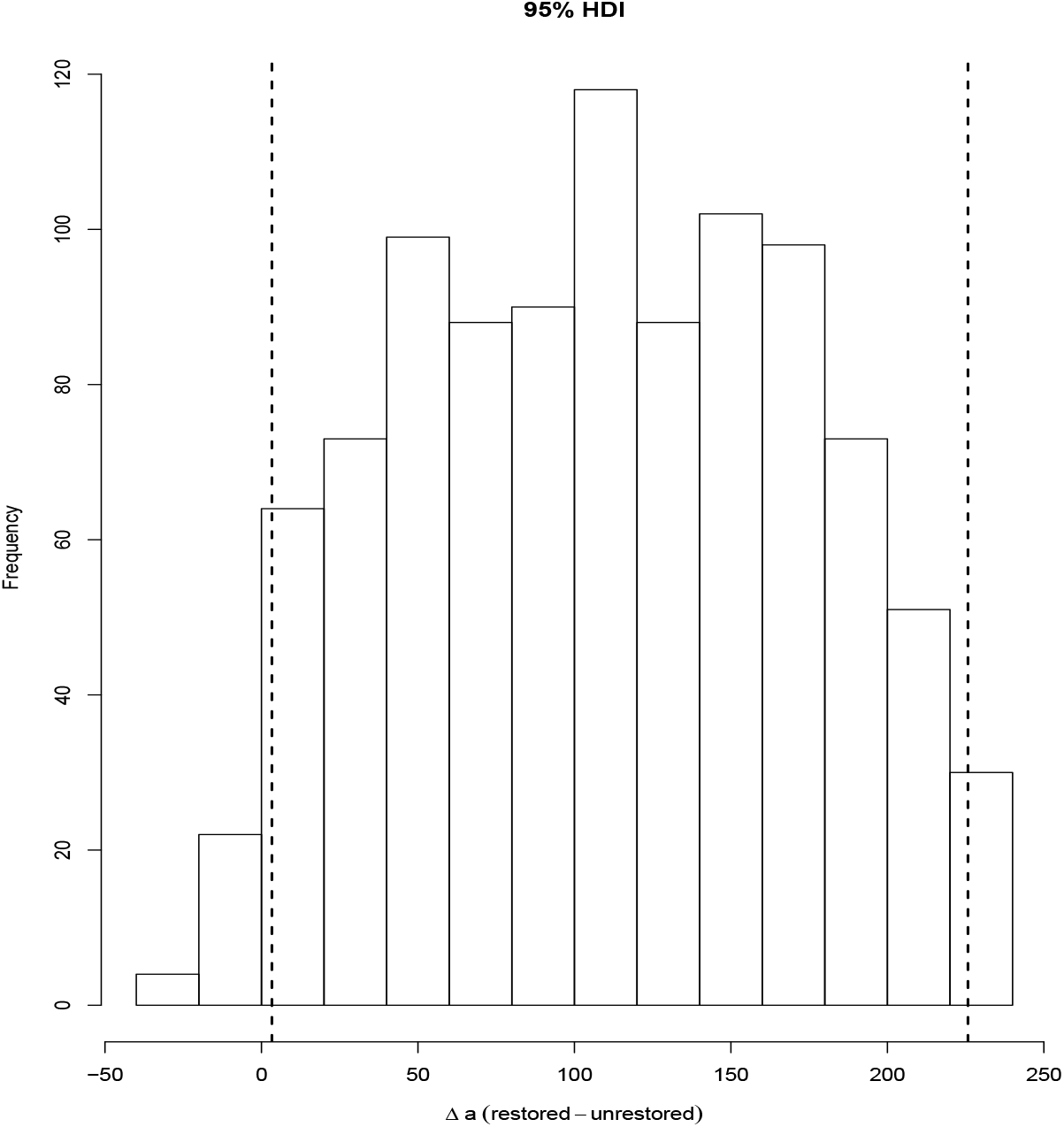
Histogram with 95% Highest Density Interval (HDI) for the habitat difference in the *a* parameter of the Equation 1 describing density-dependent immigration into restored or unrestored pools (see Fig. 3). See text for description of the derivation of the HDI. The 95% bounds (dashed lines) show *a_restored_* – *a_unrestored_* > 0, indicating restored pools have a higher immigration capacity for a given density of individuals remaining in the pool (i.e., *a_restored_* > *a_unrestored_*

The analyses for both Chinook and steelhead emigration indicated some effects of density on emigration from restored and unrestored pools (Table 3, Appendix S1: Fig. 2), consistent with movement according to habitat settlement rules. However, total number of fish marked was the strongest predictor of total number of emigrants for both species. Density was positively correlated with total emigrants for both species but there was only a significant habitat × density interaction term for steelhead, indicating a difference in the slopes of the emigrants vs. total density relationship for each habitat. These slopes showed higher density dependent emigration of steelhead from restored habitat (with 95% credible interval): 9.70 (7.98-11.42) compared with unrestored: 4.93 (2.78-7.08).

**Table 3:** Analysis of variance on a linear model showing the effects on total number of Chinook salmon (a) and steelhead (b) emigrants from restored (N = 11) and unrestored (N = 10) pools during the first 24 h after capture and marking. Significant density dependence indicated by *; total emigration was correlated with total density for both species, but density dependence via a significant habitat × density interaction term was indicated only for steelhead.

| Response | df | MS | F | *p* |
| --- | --- | --- | --- | --- |
| *a) Chinook salmon* |  |  |  |  |
| N<sub>marked</sub> | 1 | 59852 | 2980.6 | <0.0001 |
| Pool area | 1 | 161.0 | 8.02 | 0.006 |
| Total density | 1 | 99.0 | 4.92 | 0.030* |
| Habitat | 2 | 17.0 | 0.87 | 0.424 |
| Habitat × density | 1 | 17.0 | 0.88 | 0.355 |
| Resid. | 67 | 20.0 |  |  |
| *b) Steelhead* |  |  |  |  |
| N<sub>marked</sub> | 1 | 880 | 256.7 | <0.0001 |
| Pool area | 1 | 11.29 | 3.29 | 0.075 |
| Total density | 1 | 46.85 | 13.66 | 0.0005* |
| Habitat | 2 | 1.15 | 0.334 | 0.7517 |
| Habitat × density | 1 | 42.90 | 12.51 | 0.0008* |
| Resid. | 58 | 4.11 |  |  |

### Condition dependent movement

A lower coefficient of variation in condition index (CVCI) among individual Chinook salmon was negatively correlated with affinity (β) for pools with β values combined for all years and both restoration types (Pearson *r* = −0.934, P = 0.0006; Figure 5). This implies that affinity for pools (regardless of restoration status) may offer the opportunity for asset protection, resulting in reduced variability in condition among individuals. Because the regression β for restored habitat in 2016 was not significant (Table 2), it may not be valid to consider a situation with no measurable affinity for pools. Exclusion of this data point did not change the overall result (*r* = −0.874, P = 0.01). For steelhead, there was a slight negative, but non-significant correlation between the CVCI and β (Pearson *r* = −0.097, P = 0.855) and, thus, no indication that selection of pools was related to condition variability (Appendix S1: Fig. 3). Both Chinook and steelhead recaptures tended to have a lower CVCI among individuals relative to immigrants and emigrants (Appendix S1: Fig. 4) suggesting that pools may offer the opportunity for asset protection when foraging among stream habitats. Chinook recaptures where shorter and had lower mean K than both immigrants and emigrants (Appendix S1: Table 2). For steelhead both mean length and mean K were also highest in emigrants (Appendix S1: Table 3). Therefore, neither species tended to show size dominance in maintaining affinity for pools.

**Fig. 5:**
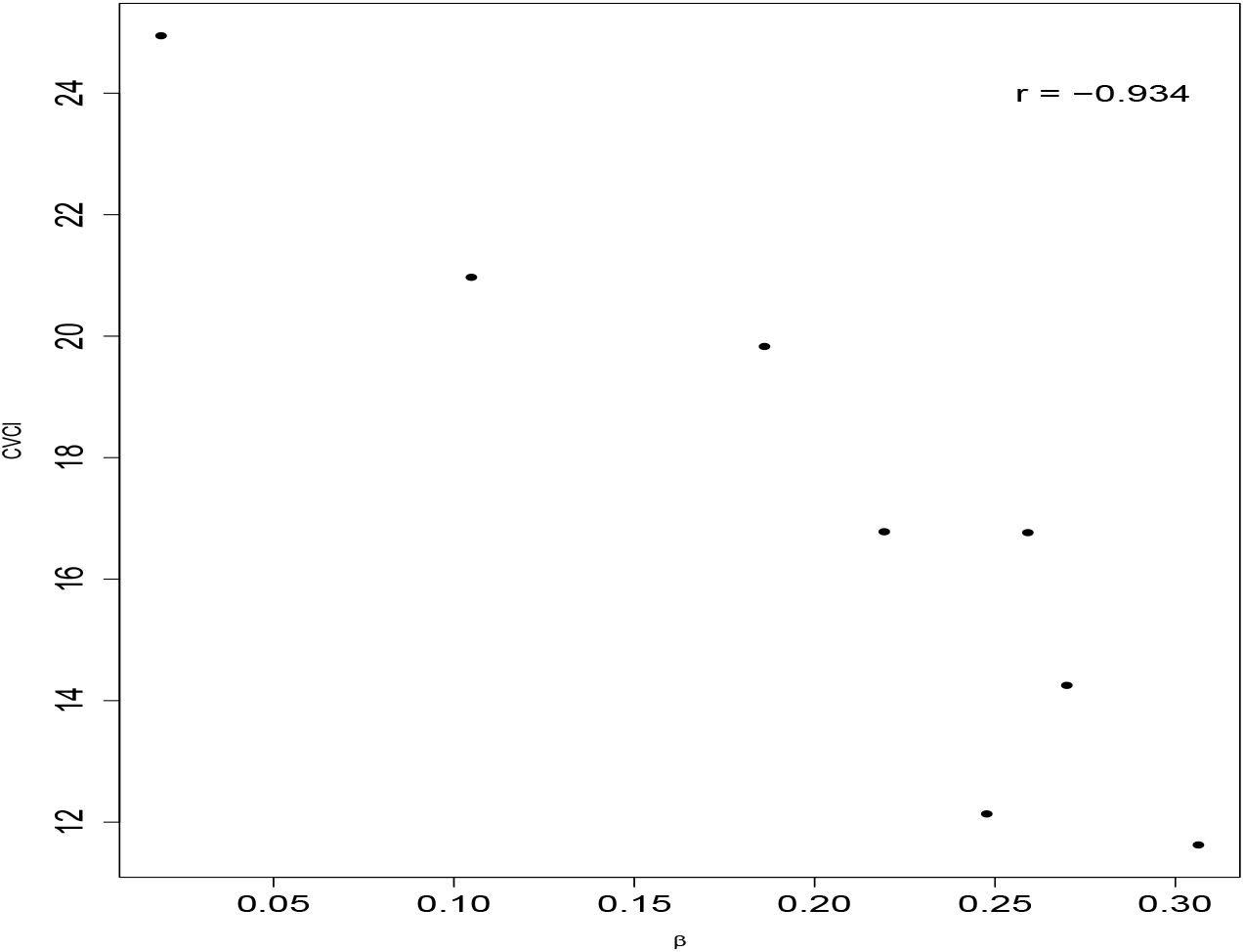
Correlation between the strength of juvenile Chinook salmon habitat affinity, indicated by the slopes (*β*) in each habitat type (taken from Table 2) and the coefficient of variation in condition index (*K*; CVCI) of recaptured individuals in each habitat type and year. When considering all years and habitats, stronger affinity for pools (regardless of restoration) was correlated with a lower CVCI (p = 0.0006).

## DISCUSSION

Habitat selection patterns indicated by mark-recapture data and additional inferences about capacity and behavior supported the positive density response to restoration identified in previous work in this study system (Polivka et al. 2015). Observations of density differences among habitats as the only metric of restoration success are often confounded by high variability among years and species, which leads to the frequent conclusion that the positive response to restoration is, at best, inconsistent (Smokorowski and Pratt 2007, Whiteway et al. 2010, Pess et al. 2012). From the standpoint of ideal free distribution theory, observed differences among restored and unrestored habitats are suggestive of quality and, thus, capacity differences (Fretwell and Lucas 1970, Houston 2008). However, under-utilization of good habitat is common (Kennedy and Gray 1993, Polivka 2005, Houston 2008), and the affinity studies presented here uncovered mechanistic details about capacity added via restoration that may not be available from distribution data. The approach described here also identified preliminary clues about asset protection (Clark 1994, Luttbeg and Sih 2010) and thus the potential fitness benefits of creating larger, better quality pools through restoration.

Although structural complexity of in-stream projects such as ELJs makes mark-recapture studies difficult when analysis depends on 100% capture, here it was possible to obtain an estimate of habitat affinity over a wide range of total fish marked. Although capture was likely less than 100% in restored pools, the resultant underestimate of density might lead to an underestimate of density on Day 1 or an underestimate of recaptures on Day 2. If Day 1 density was higher than estimated, it would have increased emigration. However, the proportion of marked fish remaining was still higher in restored pools, relative to unrestored pools that have high capture efficiency. Therefore the inferences about affinity differences between habitat types are likely conservative.

With combined data, I identified a longer term affinity trend for restored pools, but it is unclear why there was a relatively small affinity difference (*β_restored_ – β_unrestored_*) between the two habitat types for Chinook, the statistical significance of the interaction term within the best model. One explanation is that years with very strong affinity differences (e.g., 2009) contributed to the interaction term. However, this may also be an example of a statistically significant difference that is not biologically meaningful, implying no difference between restored and unrestored pools in affinity. Although this would not necessarily indicate that restoration has failed, it would show that short-term affinity, as a behavioral mechanism, did not support observations of density differences. Checking for concordance between affinity and density strengthens inferences about restoration efficacy, whereas the lack of concordance, as was apparent from some individual years in both species, indicates that single observations of density differences may be insufficient to detect fish response to restoration. Other known differences in the ecology of these two species may explain some of the variation in the results of the mark-recapture assays. Steelhead tend to be more generalist in their habitat selection pattern (Everest and Chapman 1972, Hillman et al. 1987, Young 2004) which may explain why some analyses showed selection of restored habitat by steelhead and some did not.

Most fish habitat restoration is implemented under the assumption that amelioration of some limiting physical characteristic is the key to species recovery (Roni et al. 2002, 2008, Hillman et al. 2016). Although there was generally a consistent correlation between depth and current velocity and Chinook density and habitat affinity, the models did not consistently identify a given factor for each year. Furthermore, there was almost no correlation between physical characteristics and either steelhead density or affinity/movement and, even with much more extensive sampling, habitat correlates only showed modest effects on steelhead density (Polivka et al. 2015). Thus, restoration that focuses primarily on making and measuring changes in physical habitat may neglect to address other important mechanisms and may not be suitable metrics of a realized benefit to fish. However, current velocity and temperature likely affect the energetic inputs into individual condition and movements of fish (Railsback et al. 2005). Despite little to no difference in temperature, at least between restored and unrestored pools, current velocity may still have influenced the condition data considered here. However, it is difficult to analyze the effect of current velocity on affinity at the individual pool level for the reasons described above that led to the use of GLMs to analyze pools as a group: among-pool variation in number captured causes use of individual recapture proportions to become statistically intractable (Polivka 2010).

Additional mechanistic detail about fish response to restoration came from inferences about density- and state-dependent habitat movements. Restored pools had higher capacity for immigration by Chinook across the observed density of fish remaining in pools of either habitat type. Sometimes density observations between habitat types were equivocal and this finding indicates that restoration still may result in a positive fish response. This finding is also important when density differences between habitats do exist because it suggests a non-equilibrium distribution with respect to the ideal free distribution. Immigration still occurred in both habitat types suggesting that habitat settlement was not yet complete, but was proceeding according to a qualitative difference in habitat suitability as predicted by IFD-related movements (Morris 1988) as opposed to idiosyncratic movements among habitats (Gowan et al. 1994, Rodríguez 2002). The consideration of density dependent emigration, however, suggested that pools were still possibly close to IFD equilibrium, given that Chinook emigrants left habitat types at equal rates and steelhead left restored pools at a slightly higher rate than unrestored pools. High rates of movement in these species is well-documented (Hillman et al. 1987, Railsback et al. 1999), thus additional evidence is required to confirm conclusions that this movement is influenced by habitat suitability in the context of the IFD.

This approach also has the advantage of examining the size and condition data of fish that move in and out of pools compared with fish that are recaptured. Analyses of these traits enable the generation of hypotheses about additional mechanisms of habitat selection. Emigrants tended to be longer and in higher mean condition in both species, but it is unclear why. The data identified density dependence, but larger, more robust individuals can hold territories, and might be hypothesized to maintain residency in pools (Sloat and Reeves 2014). Mean size and condition data of immigrants, emigrants and recaptures did not support territoriality as a factor that influences affinity or movement in this system. One hypothesis for the movement of these larger individuals is described as “state-dependent safety” (Luttbeg and Sih 2010). As opposed to asset protection, individuals in better condition actually take more risks and individuals in poor condition avoid risks. This results in further divergence among individuals as the more robust individuals gain further assets and those in poor condition decline further. (Luttbeg and Sih 2010, Sih et al. 2015). The larger size and condition of emigrants and consistently smaller size and lower condition of recaptures relative to emigrants might support this.

Alternatively, asset protection may drive pool use by fish as this mechanism reduces variation in condition among individuals (Clark 1994, Nonacs 2001, Conrad et al. 2011). Most of the trends observed here (Appendix S1: Fig. 4) are merely suggestive of asset protection, and with such a short-term study, it is impossible to know whether these individuals had resided in their respective pools long enough to converge in condition index. I did not quantify predation risk directly, but rather attempted to identify that signal of reduced condition variability as justification for generating hypothesis about risk-driven behavior. Similar studies also report patterns of size-dependent habitat selection suggestive of asset protection (Johnston et al. 2004, Cromwell and Kennedy 2011), but with the similar lack of distinction between predation energetic drivers of foraging in pools vs. shallower, faster flowing habitats such as riffles (Rosenfeld and Boss 2001). My study focused on restored vs. unrestored pools and thus did not compare food availability between pools of either type and riffles. However, the correlation in Chinook between the strength of affinity for pools of either habitat type and condition variation gives at least some quantitative basis worth further investigation, a two step process manifested by Polivka (2011) and Polivka et al. (2013).

Stronger evidence that reduced variability in condition among individuals is the result of asset protection can come from foraging experiments in which food availability, forager state, and risk are manipulated (Kotler et al. 2004, 2010). Observation of increased apprehension due to predators through foraging experiments with fish in the field is possible in some systems (Alofs and Polivka 2004, Polivka 2007, Balaban-Feld et al. 2019), but difficult with salmonids (Bradford and Higgins 2001). In some cases, it is relatively easy to link the trade-offs in risk to reduced condition variablity (Alofs and Polivka 2004, Polivka 2011), but other times requires the same assumptions made in this study that are based on *a priori* observations of reduced condition variability in one habitat versus another (Polivka 2011). That may remain the case here because simultaneous experimental manipulation of predation risk, food availability and forager state may be intractable in this study system and limited to lab experiments (Reinhardt and Healey 1999, Reinhardt 2002). Even if energetic optimization is a stronger driver of foraging behavior than predation risk, reduced among-individual variability might still result, for example, if individuals converge on a strategy where the marginal value of energy gained from additional foraging in faster flowing habitat no longer exceeds the cost (Girard et al. 2004). Because condition variation among individuals has implications for longer-term fitness (Clark 1994, Sinclair and Arcese 1995, Nonacs 2001, Conrad et al. 2011), and given the strong correlation between Chinook abundance and pool area in this study system (Polivka et al. 2015), creation of larger pools through restoration with ELJs may help to indirectly augment fitness for a greater number of individuals.

Site fidelity studies of this kind are applied across many taxonomic groups (Webb and Shine 1997, Sommerfeld et al. 2015) and the patterns observed can indicate the animal’s perception of habitat quality (Heap et al. 2014). To the extent that habitat quality drives habitat selection, observations of affinity may improve conclusions about the efficacy of fish habitat restoration (Bronte et al. 2007, Espinoza et al. 2011). Although mark-recapture/movement assays are not usually part of fish habitat restoration effectiveness studies (but see Freedman et al. 2016), I show here that, in a multi-year program of data collection, they may yield more robust ways of inferring benefits of restoration at the individual and, to a lesser extent, population levels. The methodology presented herein takes into account at least short term affinity which adds a mechanistic component to any observed density difference. Moreover, the density dependent immigration rate can quantify a capacity increase. Such an approach can prevent overconfidence in single observations of distribution and abundance and even indicate a benefit of restoration at times when there is no observed density difference between restored and unrestored habitat.

## Acknowledgements

Cascadia Conservation District sponsors restoration projects in the Entiat River and employees served as field assistants. I thank S. Eichler, R. Logan, A. Bushy, K. Tackman, J. West, J. Novak, K. Logan, J. Jorgensen, H. Porter, B. Forney, R. Hosman, S. Kaech, C. Skalisky, and S. Claeson for collection and curation of field data during the various years of the study. A. Bushy was supported by the American Fisheries Society Hutton Junior Fisheries Biology Scholarship. Reviews of earlier drafts and helpful input were provided by G. Dwyer, A. Rosenberger, M. Malone, S. Claeson (who designed Fig. 1 as well), R. Flitcroft, G. Pess and another anonymous reviewer.

