## Appendix S1 for "Habitat affinity and density-dependent movement as indicators of fish habitat restoration efficacy"

CARLOS M. POLIVKA*

January 16, 2020

### DENSITY-DEPENDENT IMMIGRATION - STEELHEAD

Similar to the analysis performed for Chinook (main text), I sought to examine the relationship between the density of steelhead remaining in pools (recaptures) during the assays and the number of immigrants the pool could support. As indicated, the total number of steelhead immigrants did not scale linearly with pool area. Examination of the plot of immigration ($I$) and recapture density ($R$) shows a number of points clustered around very low recapture densities and low number of immigrants (Figure S1). Indeed attempts to fit the model (Equation 1, main text) resulted in the inability to estimate the $a$ or $b$ parameters in restored pools, whereas the model was unable to estimate $\lambda$ in unrestored pools. The model thus identified no differences in density-dependent immigration among habitats for steelhead, when considered alone, consistent with a more generalist pattern of steelhead habitat use (Everest and Chapman 1972). It is possible that generalist habitat selection, combined with lower overall density in this study system (Polivka et al. 2015, 2019) would indicate that a non-linear approach to density dependent immigration that considers total density might better estimate the effect of restoration on density-dependent use of pools by steelhead.

*

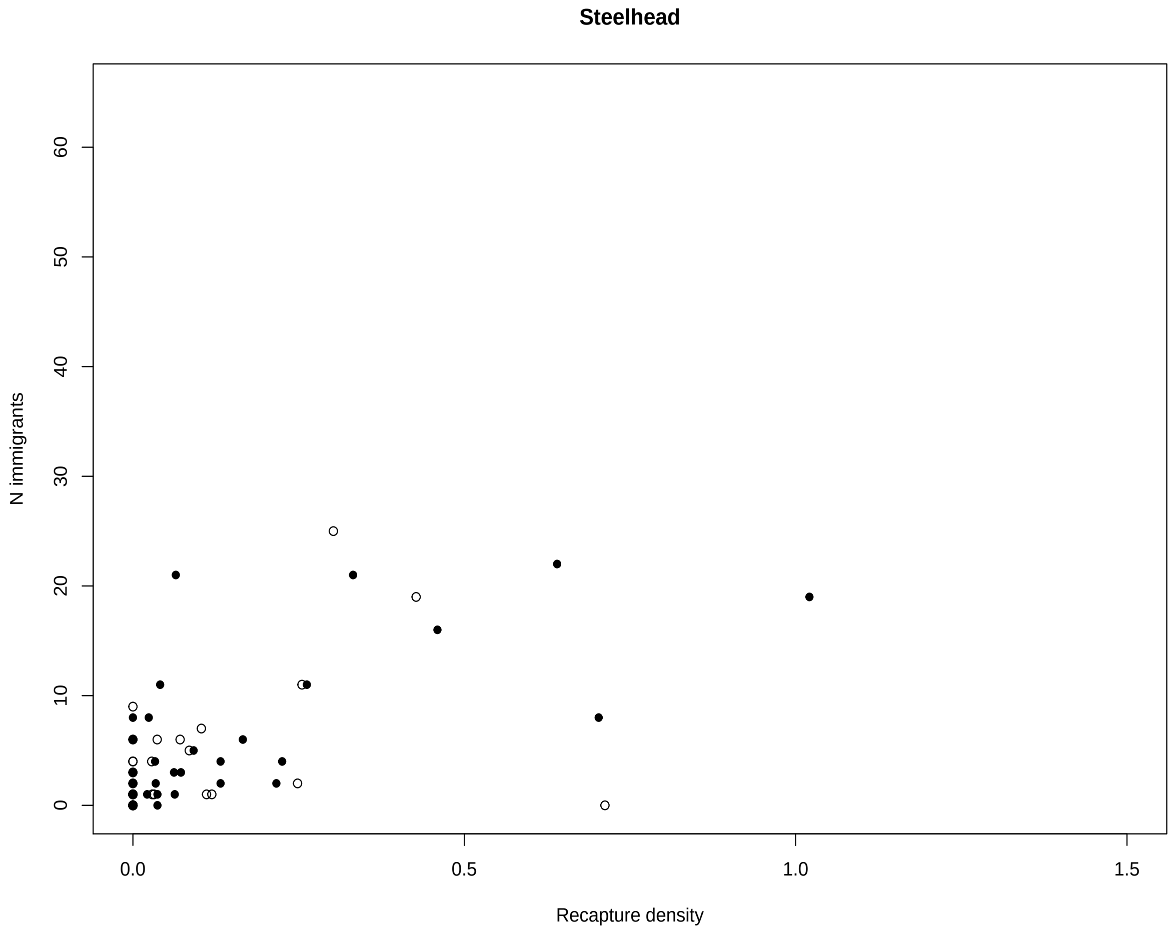

]

FIG. S 1: Immigration of young of the year steelhead into restored (solid line, filled circles) and unrestored (dashed line, open circles) pools over a 24-h period as a function of the density of fish maintaining affinity for those pools over that period (recaptures). Fit of the Ricker model curve (see main text Eq. 1) via non-linear least squares estimation of parameters showed systematic lack of fit for restored habitat and thus no comparison was made

Emigration from pools of either habitat type was expected to increase linearly with density. A generalized linear model of emigration for both Chinook and steelhead, considering total pool area, starting density and habitat type, found this linear increase (Fig. S2). Based on a habitat × density interaction term, emigration was higher in restored habitats for a given density of steelhead than in unrestored habitats, but no difference in density-dependent emigration was indicated for Chinook (see main text, Table 3).

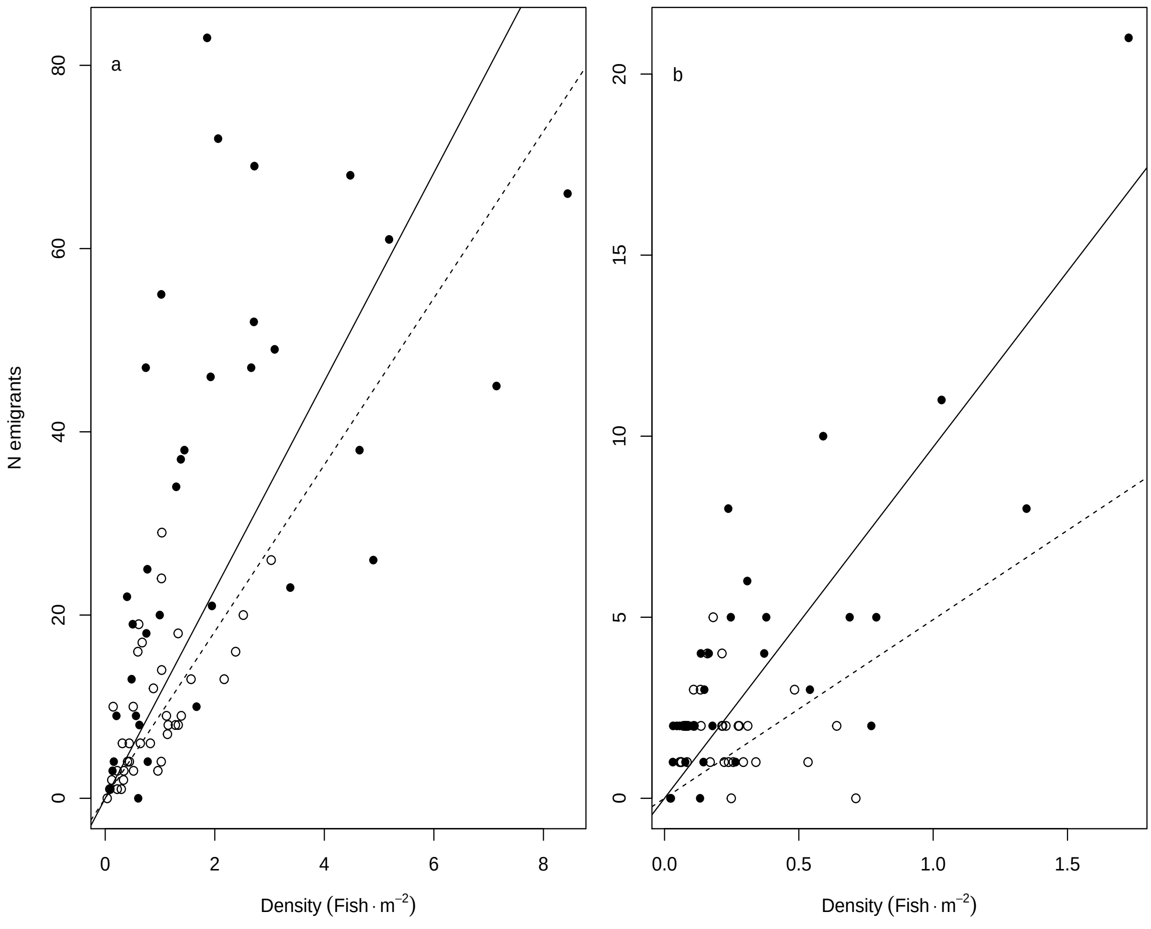

]

FIG. S 2: Number of emigrants of sub-yearling (a) Chinook and (b) steelhead into restored (solid line, filled circles) and unrestored (dashed line, open circles) pools as a function of the density of fish captured on Day 1.

### Condition-dependent Movement and Condition Variation

Under the asset protection principle (Clark 1994), trade-offs between foraging under predation risk and predator avoidance or vigilance can lead to reduced variation among individuals in condition (Nonacs 2001, van Gils et al. 2008, Luttbeg and Sih 2010). Stream pools offer habitats with the potential for predator avoidance, relative to shallower habitats with albeit higher food delivery rates (Reinhardt and Healey 1999, Railsback et al. 1999, Reinhardt 2002). Although affinity for pools as measured by generalized linear models (see main text) was associated with a decrease in variation among individuals in the Fulton Condition Index ($K$) in Chinook (main text, Fig. 5), this was not the case for steelhead (Fig. S3). Qualitative data supported the pattern observed in Chinook and also suggested that steelhead used pools in a manner consistent with asset protection principle predictions, at least in the short term. Individuals recaptured in pools 24 hrs after marking varied less among individuals in $K$ than did individuals that emigrated from the pool, or that immigrated into the pool during the interval between days (Fig. S4). It is uncertain why the two results for steelhead gave conflicting results.

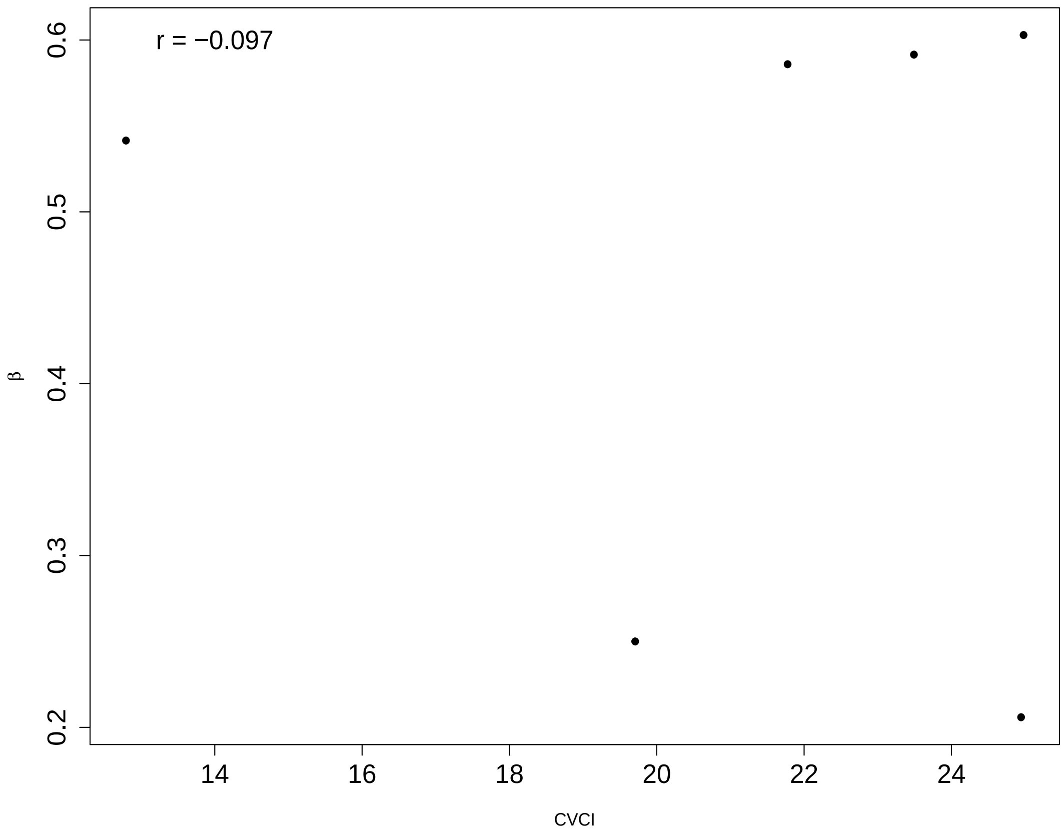

FIG. S 3: Correlation between the coefficient of variation in condition index ($K$; CVCI) of recaptured individuals in each habitat type and year (combined) and the strength of juvenile steelhead habitat affinity, indicated by the slopes ($\beta$) in each habitat type (taken from Table 2, main text). No correlation was indicated in this case. Data from unrestored habitat in 2013 and 2016 were omitted because $\beta$ values were not significantly different from zero, or were undefined.

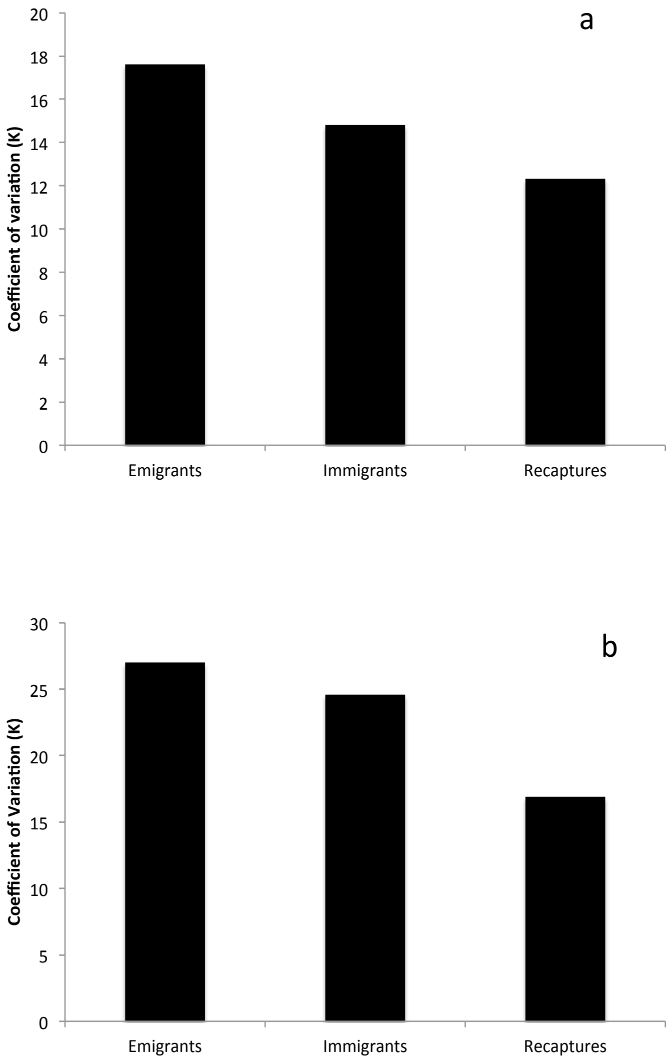

FIG. S 4: Coefficient of variation in Fulton condition index (K) for a) Chinook salmon and b) steelhead, comparing all individuals leaving a pool (both habitat types) during the 24-hr mark/recapture period ("Emigrants"), those not captured on Day 1, but found in pools on Day 2 ("Immigrants") and Day 1 fish "Recaptured" on Day 2. Recaptures tend to show the lowest variation in condition.

TABLE S 1: Total fish marked and recaptured of each species in each year of the study, restored and unrestored pools, combined. Number in parentheses is the mean per pool (both habitat types).

| Year | Chinook Marked | Chinook Rec. | Steelhead Marked | Steelhead Rec. |
|---|---|---|---|---|
| 2009 | 507 (29.1) | 133 (7.8) | 198 (11) | 107 (5.9) |
| 2012 | 580 (26.9) | 128 (5.6) | 58 (2.9) | 21 (1.1) |
| 2013 | 687 (36.2) | 219 (11.5) | 74 (4.1) | 25 (1.4) |
| 2016 | 137 (11.4) | 14 (1.2) | 16 (2) | 2 (0.25) |

TABLE S 2: Analysis of Variance on a) Length and b) Condition Index of individual sub-yearling Chinook salmon from three different movement groupings from restored and unrestored pools, combined, in 24hr mark-recapture assays. "Emigrants" = fish marked on Day 1, not recaptured on Day 2; "Immigrants" = unmarked fish captured on Day 2; "Recaptures" = fish marked on Day 1, recaptured on Day 2. Pairwise comparisons by Tukey HSD test indicated below each ANOVA result.

| Effect | df | Mean Square | F | *p* |
|---|---|---|---|---|
| *a) Length* | | | | |
| Group | 2 | 1906.3 | 17.18 | $< 0.0001$ |
| Residuals | 3563 | 110.9 | | |
| Emigrants vs. immigrants | | | | NS |
| Emigrants > recaptures | | | | $< 0.0001$ |
| Immigrants > recaptures | | | | $< 0.0001$ |
| *b) Condition Index ($K$)* | | | | |
| Group | 2 | 0.736 | 13.73 | $< 0.0001$ |
| Residuals | 3563 | 0.054 | | |
| Emigrants > immigrants | | | | $< 0.0001$ |
| Emigrants > recaptures | | | | 0.007 |
| Immigrants vs. recaptures | | | | NS |

TABLE S 3: Analysis of Variance on a) Length and b) Condition Index of individual sub-yearling steelhead from three different movement groupings from restored and unrestored pools, combined, in 24hr mark-recapture assays. “Emigrants” = fish marked on Day 1, not recaptured on Day 2; “Immigrants” = unmarked fish captured on Day 2; “Recaptures” = fish marked on Day 1, recaptured on Day 2. Pairwise comparisons by Tukey HSD test indicated below each ANOVA result.

| Effect | df | Mean Square | F | *p* |
|---|---|---|---|---|
| *a) Length* | | | | |
| Group | 2 | 2049.4 | 9.09 | 0.0001 |
| Residuals | 1301 | 225.4 | | |
| Emigrants > immigrants | | | | < 0.0001 |
| Emigrants > recaptures | | | | 0.012 |
| Immigrants vs. recaptures | | | | NS |
| *b) Condition Index ($K$)* | | | | |
| Group | 2 | 0.365 | 3.69 | 0.025 |
| Residuals | 1301 | 0.099 | | |
| Emigrants > immigrants | | | | < 0.019 |
| Emigrants vs. recaptures | | | | NS |
| Immigrants vs. recaptures | | | | NS |
